# Allele-Specific Expression and High-Throughput Reporter Assay Reveal Functional Variants in Human Brains with Alcohol Use Disorders

**DOI:** 10.1101/514992

**Authors:** Xi Rao, Kriti S. Thapa, Andy B Chen, Hai Lin, Hongyu Gao, Jill L Reiter, Katherine A. Hargreaves, Joseph Ipe, Dongbing Lai, Xiaoling Xuei, Hongmei Gu, Manav Kapoor, Sean P. Farris, Jay Tischfield, Tatiana Foroud, Alison M. Goate, Todd C Skaar, R. Dayne Mayfield, Howard J. Edenberg, Yunlong Liu

## Abstract

Transcriptome studies can identify genes whose expression differs between alcoholics and controls. To test which variants associated with alcohol use disorder (AUDs) may cause expression differences, we integrated deep RNA-seq and genome-wide association studies (GWAS) data from four postmortem brain regions of 30 AUDs subjects and 30 controls (social/non-drinkers) and analyzed allele-specific expression (ASE). We identified 90 genes with differential ASE in subjects with AUDs compared to controls. Of these, 61 genes contained 437 single nucleotide polymorphisms (SNPs) in the 3’ untranslated regions (3’UTR) with at least one heterozygote among the subjects studied. Using a modified PASSPORT-seq (parallel assessment of polymorphisms in miRNA target-sites by sequencing) assay, we identified 25 SNPs that showed affected RNA levels in a consistent manner in two neuroblastoma cell lines, SH-SY5Y and SK-N-BE(2). Many of these are in binding sites of miRNAs and RNA binding proteins, indicating that these SNPs are likely causal variants of AUD-associated differential ASE.

## INTRODUCTION

Alcohol use disorders (AUDs) are major public health problems [1-3]. Alcohol is a central nervous system depressant, and high levels of consumption over a long period may alter brain function to promote AUDs and damage the brain, in part by altering gene expression levels [4, 5]. Understanding the molecular mechanisms by which alcohol affects the brain is important and might provide clues to the causes of AUDs and ways to reverse the impact on the brain of heavy drinking.

Variations in many genes influence the risk for AUDs; however, each individual variant has only a small effect, aside from functional variants in two alcohol-metabolizing enzymes (alcohol dehydrogenase and aldehyde dehydrogenase) [1, 3, 6, 7]. In addition to genetic differences, environmental factors and interactions among the variants also affect the risk [1, 3, 6]. Genome-wide association studies (GWAS) identify regions in the genome that affect risk for complex diseases [8], but to date only a few AUD-associated loci have been unambiguously identified [7]. Most identified regions contain many variants that are inherited together (linkage disequilibrium), and associated variants may not be the causal ones.

A powerful method to study the effects of genetic variants on gene regulation is to examine Allele-Specific Expression (ASE). ASE is the difference in expression between alternative alleles and is due to regulation by *cis*-acting DNA elements, since both alleles are exposed to the same *trans*-acting environment in the cell. We integrated deep RNA-seq and single nucleotide polymorphism (SNP) genotyping data from four different brain regions of 30 subjects with AUD and 30 social/non-drinking control subjects to detect genes whose ASE differs. We then used a high-throughput reporter assay, PASSPORT-seq (parallel assessment of polymorphisms in miRNA target-sites by sequencing) [9] to systematically screen 3’ untranslated region (3’UTR) variants in those genes to determine which alter RNA levels in cells of neural origin. This assay offers a powerful tool to screen genomic variants associated with a specific phenotype to determine which affect gene regulation. In this study, we applied this assay to all the SNPs in the 3’UTR regions of the genes that demonstrated different allelic expression in any of the four brain regions between AUD and control subjects. Using this approach, we identified 25 functional SNPs that altered gene expression in the same way in both SH-SY5Y and SK-N-BE(2) cell lines. Many are located on the binding sites of miRNAs and RNA-binding proteins.

## RESULTS

### Strategy to detect differential ASE

We obtained samples from four brain regions of 60 human samples from the New South Wales Brain Tissue Resource Center (NSWBTRC) at the University of Sydney. Of these, 30 were from individuals with AUDs and 30 were from social/non-drinkers. The brain regions were the basolateral nucleus of the amygdala (BLA) that plays crucial roles in stimulus value coding and alcohol withdrawal-induced increase in anxiety [10], the central nucleus of amygdala (CE) that mediates alcohol-related behaviors and alcohol dependency [11], the nucleus accumbens (NAC) in which alcohol promotes dopamine release [12], and the superior frontal cortex (SFC) that is implicated in cognitive control and experiences neuronal cell loss after long-term alcohol abuse [13].

We performed deep RNA-seq experiments (>100 million reads per sample) from each sample (240 total samples). In addition, we obtained genotype data for all 60 subjects using the Axiom Biobank Genotyping Array (Thermo Fisher Scientific, Waltham, MA). After aligning RNA-seq reads to the human genome and imputing the SNP array results to ∼10 million SNPs, we retrieved allele counts from the RNA-seq for all SNPs for which at least one sample was heterozygous, a total of 250,007 SNPs. We focused on the ∼17,000-24,000 SNPs in each of the four brain regions that had more than 10 reads in at least 5 heterozygous samples in both AUDs and control groups. The overall strategy is summarized in Supplementary Figure 1 and Supplementary Table 1.

To examine whether the allele-specific differences vary between subjects with AUD and control, a generalized linear mixed effect model (GLMM) was implemented by comparing the number of RNA-seq reads for the reference and alternative allele at each identified SNP between the two experimental groups (AUD and control). In this model (see Materials and Methods), a negative binomial distribution was used to account for the over-dispersion of the read counts [14], and a random effect variable was introduced to model the relationship of reference and alternative read counts from the same individual. The coefficient of the interaction between allele and experimental group, *β _12_*, estimates the log2 ratio of the fraction of alternative allele reads between the control and AUD samples and was used to evaluate whether the ASE at a specific locus was significantly different between AUD and control groups.

### AUD-associated differential ASE

We identified 90 SNPs with ASE in at least one brain region that significantly differed between subjects with and without AUD (Figure 1A), evaluated by a false discovery rate (FDR) <0.05 and an estimated ratio of the fraction of alternative allele reads between the control and AUD samples larger than 2 (|*β* _12_|>1). Among these SNPs, 58 showed an increased fraction of the alternative allele in AUD samples, while 32 showed a decreased fraction. There were 26 in the BLA, 10 in the CE, 31 in the NAC, and 23 in the SFC. An example of a SNP with significant differential ASE in each brain region is shown in Figure 1B. A detailed list of these 90 SNPs is shown in Supplementary Table 2.

**Figure 1.**
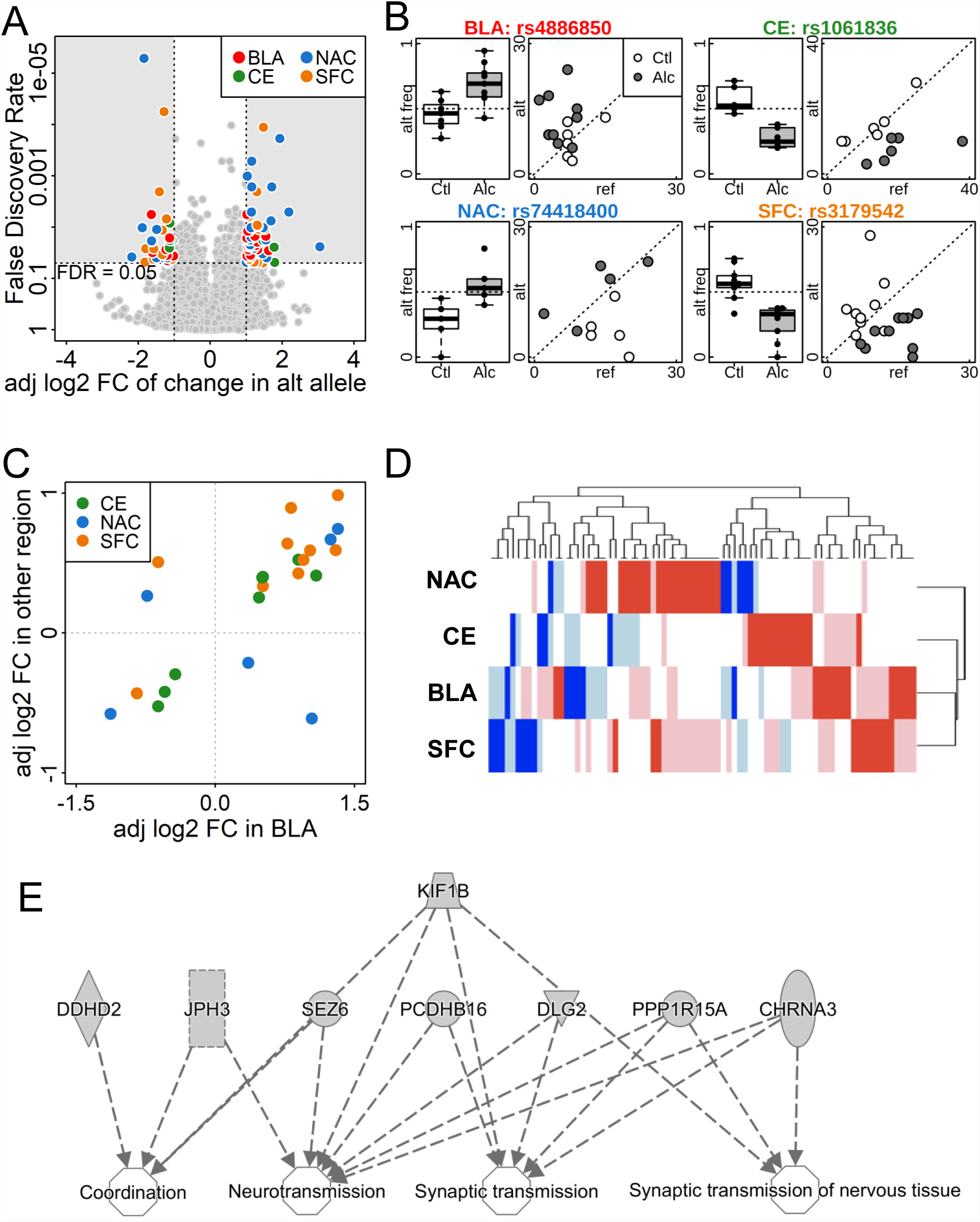
Allele-specific expression in the postmortem brain samples from AUDs subjects and social/non-drinking controls. (A) Volcano plot comparing the log2 adjusted fold change and false discovery rate of the percentage of alternative allele between AUDs subjects and controls. SNPs with FDR < 0.05 and adjusted fold change > 1 or < −1 were color-coded by brain region. BLA = basolateral nucleus of the amygdala, CE = central nucleus of the amygdala, NAC = nucleus accumbens, and SFC = superior frontal cortex; adj log2 FC = adjusted log2 fold change; alt = alternative. (B) Alternative allele frequency box plot and a scatter-plot of the number of reference and alternative reads for one significant SNP from each brain region. alt freq = alternative allele frequency; Ctl = social/non-drinking control subjects; Alc = AUDs subjects. (C) Consistency in the adjusted log2 fold change in different brain regions. SNPs with FDR < 0.05 in BLA and p < 0.05 in another brain region are plotted by adjusted fold change, color-coded by the other brain region. Of the 24 SNPs, 20 had consistent fold change direction in the two brain regions. (D) Heatmap of adjusted fold change of SNPs with FDR < 0.05 in at least one brain region shows consistency in fold change among all 4 brain regions. Dark and light colors indicate genome-wide significant (FDR < 0.05), or borderline significant (FDR > 0.05 but p < 0.05), respectively. Red and blue indicate increased and decreased percentage of alternative allele in the AUDs brain comparing to control group. (E) Ingenuity Pathway Analysis (IPA) results of genes enriched in nervous system development and function.

We examined the consistency of ASE results in the four brain regions. Among the 90 SNPs with differential ASE, many showed cross-region consistency in the coefficient of allele-alcohol interaction (*β _12_*), meaning the direction of effect was similar in multiple brain regions, although the FDR was >0.05. For example, among the 92 SNPs in BLA that had FDR<0.05 (no requirement for *β* _12_), 24 also showed significant changes in other brain regions (p<0.05), of which 20 had consistent *β _12_* direction with at least one of the other regions (Figure 1C). Similar consistency was observed in significant SNPs (FDR<0.05) in CE, NAC and SFC (Supplementary Figure 2). As shown in the heatmap (Figure 1D), among all the SNPs with FDR<0.05 in at least one brain region, a similar trend of AUD-associated allelic bias was also observed in at least one other region.

**Figure 2.**
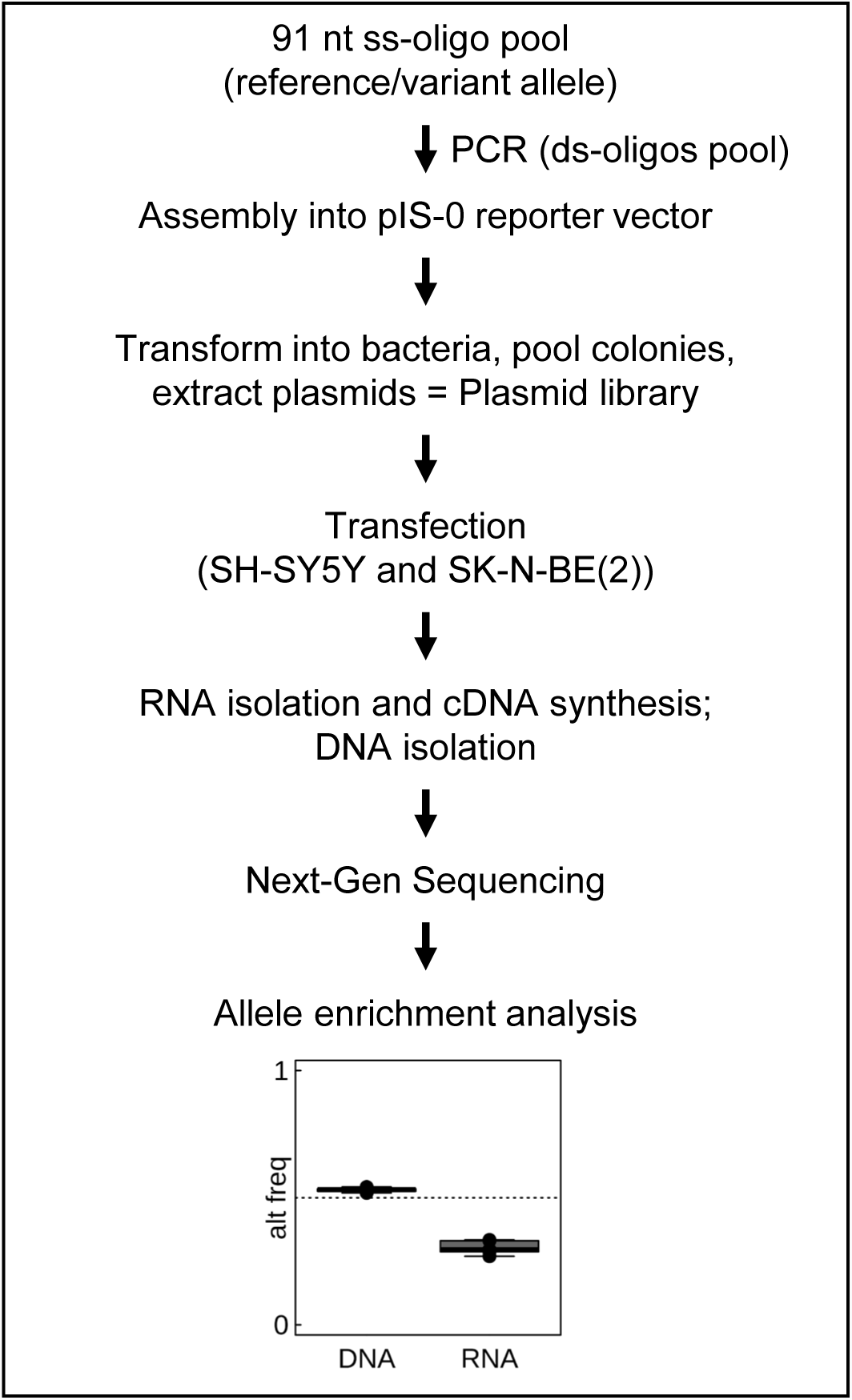
Schematic representation of the PASSPORT-seq assay. A pool of oligonucleotides flanking both alleles of 437 SNPs were synthesized (LC Sciences) and cloned in parallel into the 3’UTR of the luciferase gene in pIS-0 vector. Colonies were pooled and DNA purified. The resulting plasmid library was transiently transfected into two neuroblastoma cells lines, SH-SY5Y and SK-N-BE(2). The cDNAs were synthesized from the total RNA. The target sequences were amplified from the cDNAs and the plasmid DNA extracted from the transfected cells. The target sequences were amplified using two step-PCR with unique barcodes for each cell line and each biological replicate. Following next-generation sequencing, the reads were aligned to the reference library consisting of all the test sequences and ASE was measured for each SNPs. ss = single-stranded; ds = double-stranded.

Analyzing the genes in which the 90 ASE SNPs lie using Ingenuity Pathway Analysis showed that 17 were involved in nervous system development and function (enrichment *p-value* ranged from 8.82E-04 to 3.66E-02), including neurotransmission and coordination (Figure 1E), and 24 were involved in neurological disease (enrichment *p-value* ranged from 9.92E-04 to 3.66E-02), including movement disorders and neuromuscular disease.

### Experimental screening for functional SNPs in 3’UTR regions

Because both alleles in a heterozygote are in the same cell and exposed to the same environment, expression differences between them are due to regulation in *cis*. Among the genes with AUDs-associated differences in ASE in at least one of the four brain regions, we identified 565 SNPs in the 3’UTR regions. Among these, 437 SNPs in 61 genes had at least one heterozygote in either the AUD or control group (Supplementary Table 3). We adapted a high-throughput reporter assay, PASSPORT-seq [9], to identify 3’UTR variants that affect gene expression (RNA levels) in two neuroblastoma cell lines, SH-SY5Y and SK-N-BE(2). A schematic of the overall procedure is shown in Figure 2.

We detected reads representing reference and alternative alleles in both SH-SY5Y and SK-N-BE(2) cells in 362 (82.8%) of the 437 SNPs screened. Unique Molecular Indices (UMIs) were counted to quantify the expression of the reference and alternative alleles for each SNP of interest. A generalized linear model was applied to identify the variants whose ratio between reference and alternative allele is significantly different between the vector-expressed RNA and the plasmid DNA extracted from the cells.

As shown in Figure 3A, of the detected SNPs, 53 and 130 showed significant differences in the allelic ratio between RNA and DNA (FDR < 0.05) in SH-SY5Y and SK-N-BE(2) cells, respectively, i.e. ASE. Thirty of these showed significant differences in both cell lines, and the direction of the allele imbalance of 25 SNPs were consistent in SH-SY5Y and SK-N-BE(2): 2 from BLA, 5 from CE, 6 from NAC, and 12 from SFC (Figure 3A). Significant variants in all brain regions are shown in Supplementary Figure 3. The alternative allele frequencies and sequencing depths of four representative SNPs (one from each region) are shown in Figure 3B. Using rs1065828 in the CE as an example, the fraction of alternative allele of this SNP differ significantly between control subjects and those with AUD. The percentage of alternative UMIs in the DNA are ∼50% in both SH-SY5Y and SK-N-BE(2) cells across six biological replicates, but in the RNA it was only ∼37% in SH-SY5Y cells and ∼30% in SK-N-BE(2) cells. Since the only difference between the two alleles are the SNP loci being tested, this result indicates that this SNP led to lower expression of the alternative allele.

**Figure 3.**
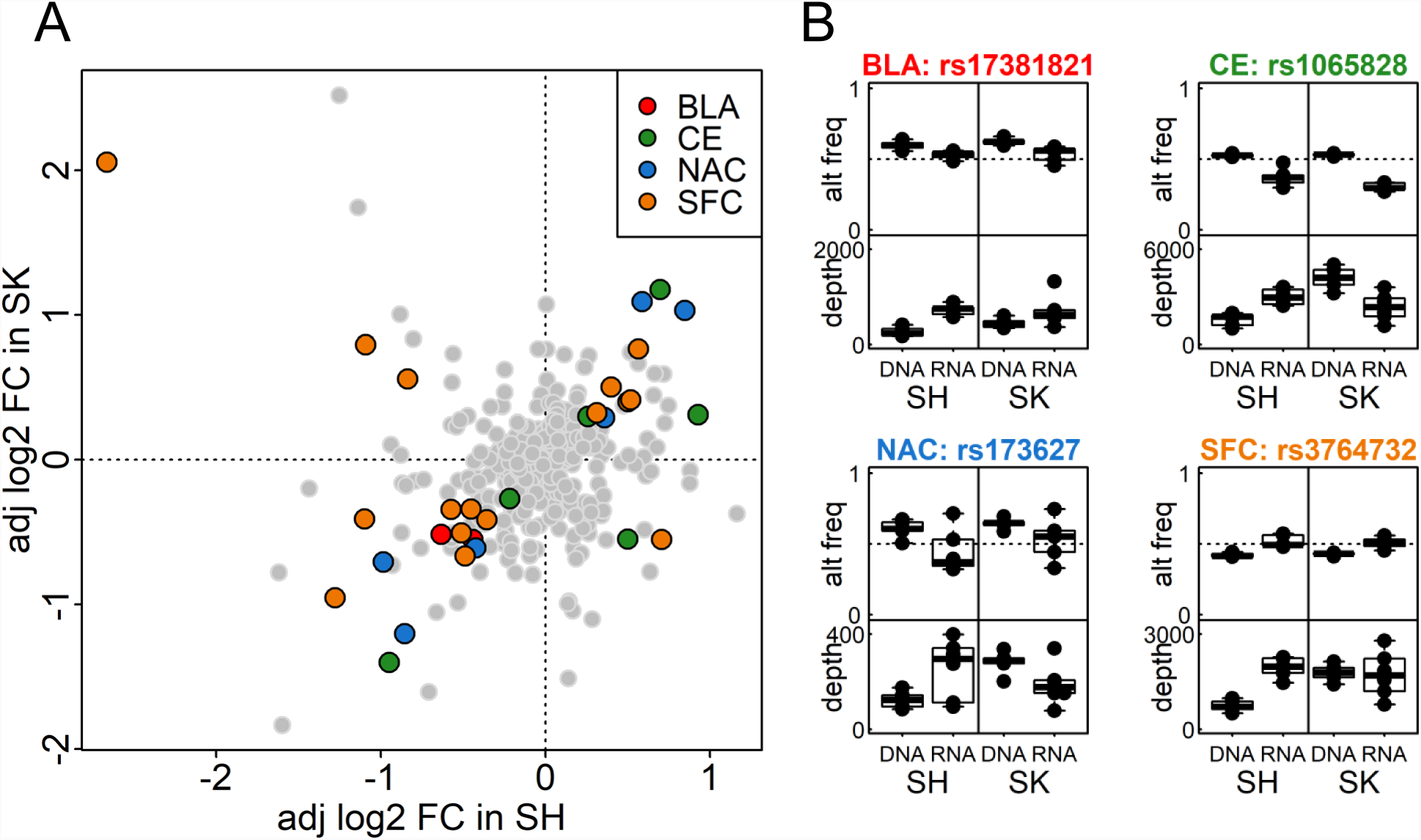
PASSPORT-seq results in SH-SY5Y and SK-N-BE(2) cell lines. (A) Plot of the adjusted log2 fold change of alternative allele frequency between RNA and DNA in SH-SY5Y and SK-N-BE(2) cell lines; a negative binomial model with random effect estimates the difference in alternative allele frequency between RNA and DNA groups (see Materials and Methods) for SNPs derived from four brain regions. SNPs with FDR < 0.05 in both cell lines were color-coded by the derived brain region. (B) Alternative allele frequency and read depth for significant SNPs derived from each brain region. SH = SH-SY5Y cell; SK = SK-N-BE(2) cell.

We annotated the 25 variants that are consistently significant in both SH-SY5Y and SK-N-BE(2) cells (Table 1). Eighteen variants were found to be in expression quantitative trait loci (eQTL) regions of their own genes based on the Genotype-Tissue Expression (GTEx) database [15]. The eQTL analysis provides additional evidence that the identified genes are, at least in part, regulated by the *cis*-acting variants. In addition, 14 SNPs coincided with the binding site of one or more RNA-binding proteins based on the ENCODE RNA-binding protein tracks, derived from RBP Immunoprecipitation-Microarray Profiling, in the UCSC Genome Browser [16]. Interestingly, Poly(A)-binding protein cytoplasmic 1 (*PABPC1*) was associated with the location of 11 SNPs, while ELAV-like RNA binding protein (*ELAVL1*) was associated with 7. TargetScan matching of miRNA seed sites at each variant location using PolymiRTS [17] found 10 SNPs that disrupt or introduce a miRNA binding site. Four of these SNPs interfere with the seed sites of 13 miRNAs that are known to be expressed in brains.

**Table 1.** Annotations for SNPs that showed significant and consistent impacts in both SH-SY5Y and SK-N-BE(2) cell lines by PASSPORT-seq

| Region | Gene | SNP | MAF | eQTL | RBP binding | Potential miRNA target |
| --- | --- | --- | --- | --- | --- | --- |
| BLA | *PEAK1* | rs17381821 | 18% | yes |  | miR-3913-3p<sup>?</sup> |
| BLA | *PNP* | rs1079375 | 32% | yes |  |  |
| CE | *HELQ* | rs4693089 | 34% | yes | PABPC1, ELAVL1 |  |
| CE | *KIF1B* | rs1065828 | 33% | yes | IGF2BP1 |  |
| CE | *KIF1B* | rs1536262 | 45% | yes | IGF2BP1 |  |
| CE | *PACRGL* | rs6812257 | 31% | yes | ELAVL1, PABPC1 |  |
| CE | *ZNF625* | rs59816741 | 1% | no |  |  |
| NAC | *LYPD5* | rs4525602 | 3% | no |  |  |
| NAC | *PCDHB16* | rs2338530 | 10% | yes | PABPC1 |  |
| NAC | *PRLR* | rs173627 | 24% | no | PABPC1 |  |
| NAC | *SLC16A4* | rs11120 | 18% | yes | ELAVL1, PABPC1 | miR-200b-3p<sup>^</sup>, miR-200c-3p<sup>?</sup>, miR-374c-5p<sup>?</sup>, **miR-429\***, miR-4330<sup>^</sup>, miR-510-5p<sup>?</sup>, miR-512-5p<sup>^</sup>, miR-655-3p<sup>^</sup>, miR-8084<sup>?</sup>, **miR-3942-3p\***, **miR-506-5p\***, miR-5100<sup>^</sup>, miR-6074<sup>?</sup>, miR-889-3p*, miR-892c-5p<sup>?</sup> |
| NAC | *SMPD4* | rs10909567 | 38% | yes | IGF2BP1 |  |
| NAC | *SYT9* | rs10732510 | 48% | no |  | miR-3925-3p<sup>^</sup>, **miR-136-5p\***, miR-4695-3p<sup>?</sup>, miR-665<sup>?</sup>, **miR-766-3p\*** |
| SFC | *CA13* | rs4617148 | 45% | yes | ELAVL1, PABPC1, IGF2BP1 | **miR-3125\***, **miR-3916\***, miR-6859-5p<sup>?</sup>, miR-6866-5p<sup>?</sup>, **miR-877-5p\***, **miR-4496\*** |
| SFC | *CA13* | rs4740047 | 42% | yes | ELAVL1, PABPC1, IGF2BP1 |  |
| SFC | *FKTN* | rs2010861 | 13% | yes | ELAVL1, PABPC1 | miR-6840-3p<sup>?</sup> |
| SFC | *LRPAP1* | rs11723071 | 20% | yes |  |  |
| SFC | *LRPAP1* | rs11729369 | 25% | yes |  |  |
| SFC | *LRPAP1* | rs13325 | 13% | no |  |  |
| SFC | *LRPAP1* | rs140653273 | 1% | no |  |  |
| SFC | *MAVS* | rs6515831 | 31% | yes |  | miR-4289<sup>?</sup> |
| SFC | *PODN* | rs899974 | 19% | no |  |  |
| SFC | *PPARA* | rs10154348 | 7% | yes | ELAVL1, PABPC1, CELF1 | miR-7155-5p<sup>?</sup> |
| SFC | *SNX18* | rs2565014 | 35% | yes | PABPC1 | miR-4655-3p<sup>?</sup><br>miR-7848-3p<sup>?</sup> |
| SFC | *UQCC1* | rs3764732 | 14% | no | PABPC1 | miR-3922-5p<sup>?</sup>, miR-513c-5p<sup>^</sup>, **miR-514b-5p\***, **miR-330-3p\***, miR-371a-5p<sup>?</sup>, **miR-371b-5p\***, miR-372-5p<sup>?</sup>, miR-373-5p<sup>^</sup>, **miR-616-5p\***, miR-7109-3p<sup>?</sup> |

\* = miRNA expressed in brain; ^ = miRNA not expressed in brain; ? = miRNA expression unknown

### Ethanol treatment changes the ASE in SNPs of interest

To evaluate the impact of alcohol treatment on the allelic imbalance of endogenous SNPs, we examined their expression in SK-N-BE(2) cells before and after treatment with two concentrations of ethanol, 10 mM and 20 mM. Among the 130 PASSPORT-seq SNPs showing ASE in SK-N-BE(2) cells (FDR<0.05 in the PASSPORT-seq assay), 17 had an average read depth per sample >15 and had at least 10% of their reads supporting the minor allele in either the ethanol-treated or untreated SK-N-BE(2) cells. Of these, we identified six whose ratios between reference and alternative alleles were significantly altered by alcohol treatment in a dose-responsive manner: rs45474901, rs2950846, rs45522239, rs45548238, rs1968676, and rs2338530. Three of these SNPs are located in the 3’UTR of *TMEM25*, two SNPs are within *PCDHB16*, and one is in *SMPD4*. The SNP with the smallest *p-value* in each gene is shown in Figure 4 (the other three are shown in Supplementary Figure 4). For 5 of these 6, the effects of acute ethanol treatment were in the opposite direction of that seen in the brains of subject with AUD. For example, the variant allele of rs45474901 (in the 3’UTR of *TMEM25)* exhibited higher expression in the PASSPORT-seq assay and in SFC of heterozygous subjects with AUDs, but lower expression after ethanol exposure in SK-N-BE(2) cells. Conversely, for rs1968676, a SNP in *SMPD4*, and rs2950846, a SNP in *PCDHB16*, the variant alleles were expressed at lower levels in AUDs subjects and in PASSPORT-seq but at higher levels after ethanol treatment.

**Figure 4.**
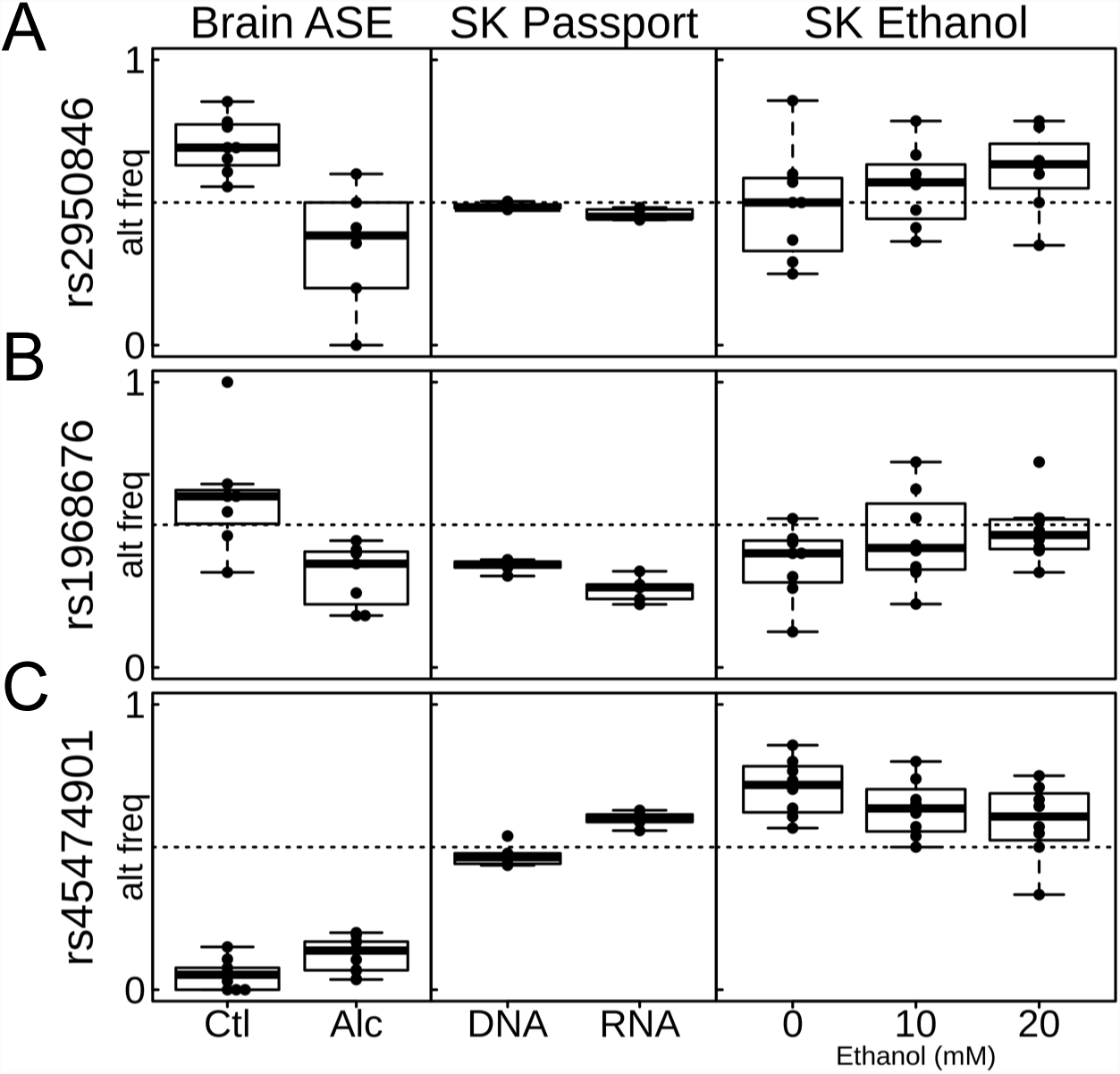
ASE, PASSPORT-seq, and Ethanol treatment results for selected SNPs. Three SNPs (rs2950846, rs1968676, rs45474901) that had alternative allele frequency significantly different in PASSPORT-seq results (FDR < 0.05) and significantly different across 0 mM, 10 mM, and 20 mM ethanol treatment dosages in SK-N-BE(2) cells (p < 0.05) were identified. The differences in the alternative allele frequency in the brains of heterozygous social/non-drinkers (Ctl) and AUDs subjects (Alc) were also shown. SK = SK-N-BE(2) cell.

## DISCUSSION

In this study, we focused on understanding the roles of *cis*-acting variants on gene regulation in AUDs. We identified 90 genes whose allele specific expression differed between AUDs and control subjects in at least one of four brain regions; many of the differences were consistent in direction across brain regions. Pathway analysis showed enrichment of these genes in pathways related to neurological disorders and neurodegenerative diseases. High-throughput screening demonstrated that 25 SNPs in the 3’UTR regions of these genes lead to ASE in both SH-SY5Y and SK-N-BE(2) cells, and many more did so in one of the cell lines.

Interestingly, the ASE of six SNPs in SK-N-BE(2) cells was altered by pharmacologically relevant concentrations of ethanol (10-20 mM); the direction of the acute alcohol response was opposite to the direction in the brain regions. Similar trends have been observed elsewhere, e.g. acute alcohol exposure favors an anti-inflammatory response and chronic alcohol consumption favors proinflammatory cytokine release [18, 19]. The SNPs with ethanol-induced changes in allelic ratio are located in the 3’UTR of the three genes, *TMEM25* (transmembrane protein 25), *SMPD4* (sphingomyelin phosphodiesterase 4), and *PCDHB16* (protocadherin beta 16). *PCDHB16* is a potential calcium-dependent cell-adhesion protein that may be involved in the establishment and maintenance of specific neuronal connections in the brain. The expression of *PCDHB16* is increased in the mouse NAC (the same region in which we found ASE) in response to cocaine [20]. *TMEM25* encodes a transmembrane protein that is expressed in multiple brain regions, including the cerebellar cortex and hippocampus, as well as in neuroblastoma and brain tumors, and may be involved in the promotion of axon growth and the regulation of cell migration [21]. *SMPD4* is also highly expressed in the brain and is important in maintaining sphingolipid metabolism in neuronal system. Sphingolipid homeostasis has been associated with the effects of alcohol on depression [22].

ASE analysis examines the transcriptional differences between two alleles and is designed to identify genes whose expression levels are influenced by the *cis*-acting elements. ASE analysis serves as a complement to eQTL analysis with higher statistical power, since the expression signals on the two alleles in the same sample serve as internal controls for each other. Despite these advantages, ASE analysis has several challenges. First, it can only focus on heterozygous loci since the analysis can only be conducted by comparing the expression signals of two different alleles. Second, ASE analysis from RNA-seq data is subject to technical challenges due to the differences in our ability to align sequence reads to two alleles. Normally, most alignment algorithms more readily align the sequencing reads with the reference sequence, which can lead to depressed expression signals for the alternative allele. Fortunately, this type of potential bias can be avoided by comparing the allelic imbalances between two experimental conditions since the biases due to alignment algorithms will be the same in both. Thus, we focused our analysis on the differences in allelic imbalance between subjects with and without AUDs. This strategy also allows us to focus our analysis on the genetic variants that are associated with AUDs.

In this study, we used a high-throughput reporter assay, PASSPORT-seq [9], to screen for 3’UTR variants that lead to gene expression changes. We modified the original protocol to improve the quality of the data. First, we introduced UMIs during the reverse transcription of the mRNA and in the first PCR of the plasmid DNA used as a control to overcome potential PCR biases during the library construction. Second, we added staggered sequences to the PCR primers to reduce problems associated with low sequencing complexity at the beginning of the reads (Supplementary materials). Differences in ASE between subjects with and without AUDs could be preexisting or the result of decades of excessive alcohol consumption. The functional SNPs, however, were detected in cultured cells in the absence of ethanol and therefore likely affect expression the brain even absent ethanol exposure. A limitation is that we only screened 3’UTR variants; variants in other regulatory regions, such as enhancers and promoters, also play important roles in *cis*-acting regulation.

In summary, this study identified a subset of genes that show ASE differentially between subjects with and without AUDs; the differences might preexist and affect the risk for AUDs or might result from alcohol-induced neurological damage. Within those genes, we identified SNPs that affect gene expression levels in neuronal cells, and are likely to affect expression in the brain. By performing PASSPORT-seq, our analysis goes one step deeper in understanding the genetic mechanisms of AUD. We believe similar assays should be routinely implemented to screen for functional genetic variants suggested by statistical genetics analysis, such as from genome-wide association or next generation sequencing-based analysis.

## MATERIALS AND METHODS

### Human brain tissues

The human brain tissues of four brain regions were obtained from postmortem individuals from the New South Wales Brain Tissue Resource Center (NSWBTRC) at the University of Sydney. The 60 individuals were composed of 30 subjects with AUDs and 30 social/non-drinkers. Alcohol dependent diagnoses of the 30 AUDs subjects met the APA (American Psychiatric Association) DSM-IV criteria, and were confirmed by physician interviews, review of hospital medical records, questionnaires to next-of-kin, and from pathology, radiology, and neuropsychology reports. The control group includes 30 social/non-drinker samples, which were matched with the 30 AUDs subjects based on age, sex, postmortem interval, pH of tissue, disease classification, and cause of death. Additional criteria include greater than 18 years of age, no head injury at time of death, lack of developmental disorder, no recent cerebral stroke, no history of other psychiatric or neurological disorders, no history of intravenous or polydrug abuse, negative screen for AIDS and hepatitis B/C, and postmortem interval within 48 hours.

### Data processing of RNA-seq and GWAS SNP array of human brain tissues

The mRNA was extracted from the brain tissue using Qiagen RNeasy kit (Qiagen, Germantown, MD). The RNA-seq samples were prepared using the TruSeq RNA Library Pre Kit v2 and sequenced on the Illumina HiSeq 2000. Paired-end libraries with an average insert size of 180 bp were obtained.

Raw reads were aligned to human genome 19 (hg19) using STAR aligner (version 2.5.3.a). We used FastQC (https://www.bioinformatics.babraham.ac.uk/projects/fastqc/)to evaluate RNA sequence quality. Picard “MarkDuplicates” option was used to flag and remove duplicate reads.

DNA was obtained also from the brain tissue. The Axiom Biobank Genotyping Array (ThermoFisher Scientific, Grand Island, NY, USA) was used for genotyping. Monomorphic variants, variants with call rate <=0.98 and Hardy-Weinberg Equilibrium P<10^-5^; and samples with call rate <0.90 were removed using PLINK [23]. Phasing was done using SHPAEIT2 [24] then IMPUTE2 [25] was used for imputation with 1,000 Genomes Phase 1 integrated panel excluding singleton variants as the reference for imputation. Variants with imputation score >=0.8 and estimated MAF >=0.5 were included in analysis.

### Allele-specific expression analysis

GATK ASEReadCounter was used to obtain reference and alternative allele counts at the exonic SNPs. We only analyzed SNPs with at least five heterozygotes in both control and AUDs groups, all of which had more than 10 reads, to ensure that we had enough samples with reliable read counts for analysis.

We used a generalized linear mixed effect model (GLMM) to model the number of RNA-seq reads for each allele based on its allelic type (reference or alternative allele), study group (AUDs and social/non-drinkers), and the interaction terms between the two variables. A random variable was used to account for the reads from the two alleles within the same individual. To adjust for the overdispersion effects of the RNA-seq reads, a negative binomial distribution was used in the GLMM model.

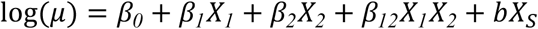

where *μ* is the expected number of sequencing reads for one allele (reference or alternative) in one specific subject, X_1_ is the allele type (0: reference allele, and 1: alternative allele), X_2_ is the subject study group (0: control group, and 1: AUDs group), and X_S_ is the subject ID. In this model, *β _0_, β _1_, β _2_*, and *β _12_* are coefficients of fixed effects, while b is coefficient for random effect that models the differences between subjects. Our null hypothesis (H_0_) is: *β _12_*= 0. Rejecting the null hypothesis indicates that the allelic imbalance differs in AUDs and control group. False discovery rate was calculated using the Benjamini-Hochberg procedure [26]. Ingenuity Pathway Analysis (IPA) is used for deriving the pathways of the genes with AUDs-associated allelic differences.

### High-throughput reporter (PASSPORT-seq) assay

The PASSPORT-seq assay was conducted as previously described [9] with some modifications. Briefly, the procedure includes the following steps: oligonucleotide synthesis, plasmid library construction, transfection, DNA/RNA extraction, and sequencing. The oligonucleotide pool was synthesized using Oligomix^®^ by LC Sciences (Houston, TX) and included 874 DNA oligos targeting both alleles of 437 SNPs in the 3’UTR of genes that showed differences in allelic ratio between control and AUDs groups. These were cloned in parallel into the 3’UTR of the luciferase gene in the pIS-0 vector (12178, Addgene, Cambridge, MA) [27] and DNA extracted and purified. The resulting plasmid pool was transfected into two neuroblastoma cell lines, SH-SY5Y and SK-N-BE(2). Each transfection was repeated six independent times. After 42 hours of transfection, both plasmid DNA and RNA were extracted from the cells. cDNA was synthesized and sequences containing the SNPs were amplified using PCR primers that also contained sample barcodes, unique molecular indices and Illumina sequencing adapters. The resulting PCR products were sequenced using one lane of Illumina HiSeq 4000 using a 75 bp paired-end protocol. The detailed experimental protocol is provided in the supplementary materials.

### Analysis of the high-throughput reporter assay

FASTQ files were demultiplexed using “cutadapt”[28] based upon barcodes identifying DNA source and replicate number, and the barcodes and adapter sequences were trimmed from each read. The UMI was then trimmed and stored in the read name using “umi_tools” [29]. Using “bwa mem” [30], remaining reads were mapped to reference sequence defined by the reference and alternative sequence for each SNP of interest. Finally, “umi_tools” was used to count the number of UMI-unique reads for each sequence.

To identify the functional 3’UTR SNPs, a generalized linear model was used to modeling differences between plasmid DNA and RNA extracted from the transformed cells in the ratio of of UMI counts supporting reference and alternative alleles. Similar to ASE analysis, a negative binomial distribution was used to account for the over-dispersion effects on the count data.

### Annotation on the functional 3’UTR SNPs selected by PASSPORT-seq assay

For each SNP, eQTLs (from GTEx), RNA-binding protein binding sites, and miRNA target sites were annotated. The eQTL information is retrieved from the Genotype-Tissue Expression (GTEx) database [31] to find SNPs that are eQTLs of the same gene. The UCSC Genome Browser database [32] was used to annotate whether a specific SNP is located in the binding site of an RNA-binding protein using the ENCODE RNA Binding Protein track [16]. The Polymirts database [17] was used to predict the potential miRNAs whose binding can be altered by a candidate SNP, where the binding of an miRNA on a specific target sequence is evaluated by TargetScan [33]. GTEx database [31] was used to determine whether an miRNA is expressed in brain.

### Ethanol treatment of cells

To evaluate the potential effect of alcohol on the function of the 3’UTR SNPs, untransfected SK-N-BE(2) cells were grown to confluence in a 9.5 cm^2^ Corning^®^ CellBIND^®^ 6-well dish (Corning, Corning, NY), then treated with physiological concentrations of ethanol (10 mM or 20 mM) or left untreated. After 24-hr and 42-hr post ethanol treatment, cells were harvested for RNA isolation, library preparation and RNA sequencing. For the 42-hr treatment condition, media was replaced with fresh media with or without ethanol in both ethanol treated and control cells respectively at 24-hr. Four independent biological replicates were conducted for each condition. The mRNA was extracted from the cells using Qiagen RNeasy kit (Qiagen, Germantown, MD). The RNA-seq samples were prepared using the TruSeq RNA Library Pre Kit v2 (Illumina, Inc., San Diego, CA) and sequenced on the Illumina HiSeq 4000 using 2×75bp paired-end configuration. A *GLMM* model was used to identify the variants whose allelic frequencies are altered by ethanol treatment (see supplementary materials).

## Supplementary Figures

**Supplementary Figure 1.** Workflow for identifying SNPs in the 3’UTRs of genes with differential allele-specific expression in heavy drinkers compared with non-drinkers. (A) Heterozygous SNPs are identified in the RNA-sequencing data from brain tissue in social/non-drinkers and heavy drinkers. Differential allele-specific expression is detected using a negative binomial model with random effects. (B) Genes with differential ASE are identified and 3’UTR SNPs from those genes are identified. Passport-seq screening of these SNPs help to identify candidate SNPs for further validation. Het. = heterozygous.

**Supplementary Figure 2.** Consistency of differential ASE results between brain regions. For each significant (FDR < 0.05) gene in CE that is significant (p < 0.05) in another brain region (BLA, NAC, SFC), the adjusted log2 fold change was plotted. This was repeated for each brain region. Most genes had the same direction of change in each pair of brain regions. (BLA = basolateral nucleus of the amygdala, CE = central nucleus of the amygdala, NAC = nucleus accumbens, and SFC = superior frontal cortex).

**Supplementary Figure 3.** Passport-seq results for other brain regions (BLA, CE, NAC, and SFC). Significantly differentially expressed genes in SH-SY5Y and SK-N-BE(2) cells are color-coded to denote significance. SH = SH-SY5Y cell; SK = SK-N-BE(2) cell.

**Supplementary Figure 4.** ASE, PASSPORT-seq, and Ethanol treatment results for additional ethanol SNPs with alternative allele expression sensitive to ethanol treatment. Three additional SNPs (rs45522239, rs45548238, rs2338530) that had alternative allele frequency significantly different in PASSPORT-seq results (FDR < 0.05) and significantly different across 0 mM, 10 mM, and 20 mM ethanol treatment dosages in SK-N-BE(2) cells (p < 0.05) were identified. The differences in the alternative allele frequency in the brains of heterozygous social/non-drinkers (Ctl) and AUDs subjects (Alc) were also shown. SK = SK-N-BE(2) cell.

## TABLE LEGENDS

**Table 1**. **Annotations for SNPs that showed significant and consistent impacts in both SH-SY5Y and SK-N-BE(2) cell lines by PASSPORT-seq.** For each SNP, the minor allele frequency (MAF), whether the SNP had a significant eQTL for its gene in GTEx, RNA-binding proteins (RBP) whose binding sites overlapped the SNP, and miRNAs whose binding potential was changed by the alternative allele.

## ACKNOWLEDGEMENTS

We thank our laboratories members for critical discussions and the Center for Medical Genomics facility at Indiana University School of Medicine for technical support. This work was supported by NIH grant U10AA008401, from the National Institute on Alcohol Abuse and Alcoholism (NIAAA), as part of the Collaborative Study on the Genetics of Alcoholism (COGA), and U01AA020926. The COGA, Principal Investigators B. Porjesz, V. Hesselbrock, H. Edenberg, L. Bierut, includes 10 different centers: University of Connecticut (V. Hesselbrock); Indiana University (H.J. Edenberg, J. Nurnberger Jr., T. Foroud); University of Iowa (S. Kuperman, J. Kramer); SUNY Downstate (B. Porjesz); Washington University in St. Louis (L. Bierut, A. Goate, J. Rice, K. Bucholz); University of California at San Diego (M. Schuckit); Rutgers University (J. Tischfield); Southwest Foundation (L. Almasy); Howard University (R. Taylor); and Virginia Commonwealth University (D. Dick). Other COGA collaborators include: L. Bauer (University of Connecticut); D. Koller, S. O’Connor, L. Wetherill, X. Xuei (Indiana University); Grace Chan (University of Iowa); N. Manz, M. Rangaswamy (SUNY Downstate); A. Hinrichs, J. Rohrbaugh, J-C Wang (Washington University in St. Louis); A. Brooks (Rutgers University); and F. Aliev (Virginia Commonwealth University). A. Parsian and M. Reilly are the NIAAA Staff Collaborators. We continue to be inspired by our memories of Henri Begleiter and Theodore Reich, founding PI and Co-PI of COGA, and also owe a debt of gratitude to other past organizers of COGA, including Ting-Kai Li (currently a consultant with COGA), P. Michael Conneally, Raymond Crowe, and Wendy Reich, for their critical contributions.

