## Supplemental Information for "Allele-Specific Expression and High-Throughput Reporter Assay Reveal Functional Variants in Human Brains with Alcohol Use Disorders"

**Revised PASSPORT-seq assay**

The PASSPORT-seq assay was carried out essentially as previously described [1] but with some modifications, described below, to increase accuracy.

*Oligonucleotides*

874 single-stranded DNA oligos (437 reference and 437 variants) were commercially synthesized in parallel as a single pool (Oligomix®, LC Sciences, Houston, TX). Each oligo was 91 nt long with the SNP in the center of the sequence and flanked by 25 nt on each side identical to the genomic context of the SNP, and with complementary regions from the vector on both ends (5’ end GCCGTGTAATTCTAGGAGCTC; 3’ end CGTTCTAGAGTCGGGGCGG) to allow cloning by the NEBuilder® HiFi DNA assembly reaction (New England Biolabs (NEB), Ipswich, MA). A list of the oligonucleotides is in **Supplementary Table 4**. About seven nanograms of the oligonucleotide pool was amplified by PCR using 0.5 µM of primers HJ7027 (5’ GCCGTGTAATTCTAGGAGCTC 3’) and HJ7028 (5’ GCCCCGACTCTAGAACG 3’) and Invitrogen Platinum SuperFi DNA Polymerase (Thermo Fisher Scientific, Waltham, MA) in a 20 µL reaction. The PCR conditions were: 98 °C for 30 s, 25 cycles of 98 °C for 5 s, 65 °C for 10 s and 72 °C for 10 s, with a final extension at 72 °C for 5 min. The size of the PCR product was verified on a 2% agarose gel.

*PASSPORT-seq library construction*

The pIS-0 vector (12178, Addgene, Cambridge, MA) [2] (**Figure S1**) was linearized with *Sac*I-HF® and *Bmt*I-HF® restriction enzymes (NEB). The linearized plasmid was separated on a 0.8% agarose gel, excised and purified using QIAquick gel extraction kit (Qiagen, Germantown, MD). 50 ng of the linearized plasmid and 2 µL of the amplified double-stranded oligonucleotide pool were assembled using NEBuilder® HiFi DNA assembly kit (NEB) following manufacturer’s instructions. Two µL of the undiluted assembled product were transformed into 50 µL chemically competent NEB® 5-alpha Competent *E. coli* cells (NEB) per manufacturer’s protocol. Three independent plasmid assemblies were performed with a total of six transformations (four transformations from the first assembly and one transformation each from the second and third assemblies). The cells from each transformation were plated onto five 100 mm LB-agar plates containing 100 µg/ml ampicillin. Following incubation overnight at 37°C, the colonies were dislodged from the plates by adding 1 mL sterile water per plate and collected using L-shaped cell spreaders (Thermo Fisher Scientific). A total of ~20,000 colonies were collected. Plasmid DNAs from each plate were isolated using PureLink® HiPure Plasmid Filter Midiprep Kit (Thermo Fisher Scientific) or QIAprep Spin Miniprep Kit (Qiagen). The DNAs were pooled to create a single plasmid library. DNA concentrations were measured using the Qubit™ double stranded DNA Broad Range assay kit (Thermo Fisher Scientific) and a NanoDrop^TM^ 2000 spectrophotometer (Thermo Fisher Scientific).

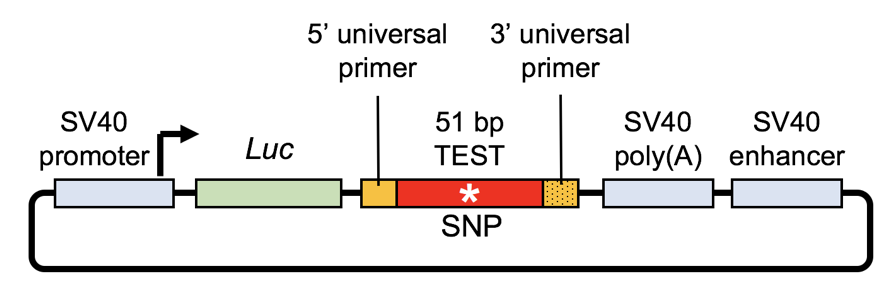

**Figure S1. Map of pIS-0 reporter vector used in PASSPORT-seq assay.** Test fragments (51 bp with the variant at position 25, flanked by 5’ and 3’ universal primers) were cloned downstream of the firefly luciferase (*Luc+)* gene within the 3’UTR.

*Sanger sequencing*

Plasmids from 60 individual colonies were isolated using QIAprep® Spin miniprep columns (Qiagen) and sequenced using primer HJ7209 (5’ GTGGTTTGTCCAAACTCATC 3’) by Sanger sequencing (Genewiz, South Plainfield, NJ) to determine the representation of the constructs.

*Culturing and transfection of SH-SY5Y and SK-N-BE(2) cells*

Two neuroblastoma cells lines, SH-SY5Y (CRl-2266, ATCC, Manassas, VA) and SK-N-BE(2) (CRL-2271, ATCC) were cultured in a 1:1 mixture of EMEM (ATCC) and F12K medium (10025-CV, Thermo Fisher Scientific) with 10% (vol/vol) fetal bovine serum (ATCC) and 1% penicillin and streptomycin. Cells were plated at a density of 9 x 10^5^ per well in a 9.5 cm^2^ Corning® CellBIND® 6-well dishes (Corning, Corning, NY). 24 h after seeding (~70-80% confluence), the medium was changed to a similar medium but without antibiotics. Cells were transfected with 1.5 µg of plasmid library using 6 µL of X-tremeGENE HP DNA transfection reagent (4:1 reagent-to-DNA ratio; Millipore Sigma, St. Louis, MO) in 200 µL Opti-MEM reduced serum medium (Thermo Fisher Scientific). At 42 h post transfection, cells were washed once with 2 mL cold 1X phosphate-buffered saline and resuspended in 700 µL of QIAzol Lysis Reagent (Qiagen). Cells were harvested and combined from two wells of a 6-well dish for RNA and DNA isolation.

*RNA and DNA isolation and cDNA synthesis*

Both plasmid DNA and total RNA were isolated from transiently transfected cells using miRNeasy mini kit (Qiagen) per manufacturer’s instructions. After the upper aqueous phase was collected for RNA isolation, DNA was extracted from the interphase and the lower phenol-chloroform phase using ethanol precipitation and resuspended in 100 µL of 8 mM NaOH. The plasmid DNA was then purified using QIAprep® Spin miniprep columns (Qiagen). RNA was purified from the upper aqueous phase using miRNeasy mini kit columns (Qiagen), with on-column DNase treatment. The RNA quantity was measured using a NanoDrop^TM^ 2000 spectrophotometer (Thermo Fisher Scientific), and 1 µg of total RNA was used for cDNA synthesis using QuantiTech Reverse Transcription kit (Qiagen) with one of the four reporter-RNA specific primers (HJ7210-HJ7213, **Table S1**) in a 20 µL reaction.

*Generating barcoded sequencing libraries from cDNAs and plasmid DNAs*

For generating sequencing libraries, PCR primers containing the flanking sequences from the vector (universal primer, 22 bp RP or FP), a set of unique molecular indices (UMIs, 10 random nucleotides, to reduce PCR bias) and a sample barcode (BC, 9-12 bp with staggered sequences, to reduce low sequencing complexity problems) were used (**Table S1, Figure S2)**. Partial Illumina adapter sequences (20 bp R1 or 21 bp R2) were also included on the primers to facilitate the library preparation (**Table S1, Figure S2**). The first PCR was performed using 2 µL of unpurified cDNA and adding 0.1 µM of one of the primers (HJ7214-HJ7219; **Table S1**; the primer carried over from cDNA synthesis was also present) in a 20 µL reaction using the following conditions: 98 °C for 30 s, 10 cycles of 98 °C for 5 s, 65 °C for 10 s, with a final extension at 72 °C for 5 min.

**Table S1. Primers used for barcoding cDNAs and plasmid DNAs**

| **Name** | **Description** | **Sequence** |
| --- | --- | --- |
| HJ7210 | 1st strand-cDNA | ACA CGA CGC TCT TCC GAT CTN TAG GCT CTN NNN NNN NNN CGG CCG CCC CGA CTC TAG AAC G |
| HJ7211 | 1st strand-cDNA | ACA CGA CGC TCT TCC GAT CTN NGA AGA CTG NNN NNN NNN NCG GCC GCC CCG ACT CTA GAA CG |
| HJ7212 | 1st strand-cDNA | ACA CGA CGC TCT TCC GAT CTN NNC ATT GCA CNN NNN NNN NNC GGC CGC CCC GAC TCT AGA ACG |
| HJ7213 | 1st strand-cDNA | ACA CGA CGC TCT TCC GAT CTN CGG AAG AAN NNN NNN NNN CGG CCG CCC CGA CTC TAG AAC G |
| HJ7214 | 2nd strand | CAG ACG TGT GCT CTT CCG ATC NAA TCC AGG CGC CGT GTA ATT CTA GGA GCT C |
| HJ7215 | 2nd strand | CAG ACG TGT GCT CTT CCG ATC NNT GAG GAG ACG CCG TGT AAT TCT AGG AGC TC |
| HJ7216 | 2nd strand | CAG ACG TGT GCT CTT CCG ATC NNN GAC TTG GAC GCC GTG TAA TTC TAG GAG CTC |
| HJ7217 | 2nd strand | CAG ACG TGT GCT CTT CCG ATC NTC TCA CCA CGC CGT GTA ATT CTA GGA GCT C |
| HJ7218 | 2nd strand | CAG ACG TGT GCT CTT CCG ATC NNG TGC GTT ACG CCG TGT AAT TCT AGG AGC TC |
| HJ7219 | 2nd strand | CAG ACG TGT GCT CTT CCG ATC NNN TCA TCG AGC GCC GTG TAA TTC TAG GAG CTC |
| HJ7220 | Passport-Seq-PCR | ACA CGA CGC TCT TCC GAT CT |
| HJ7221 | Passport-Seq-PCR | CAG ACG TGT GCT CTT CCG ATC |

The PCR products were purified using HP PCR product purification kit (Millipore Sigma) and eluted in 20 µL TE (10 mM Tris-HCl, 1 mM EDTA, pH 8.0). 9.2 µL of the products from the initial PCR was then used as template for the second PCR using 0.2 µM primers HJ7220 (5’ ACA CGA CGC TCT TCC GAT CT 3’) and HJ7221 (5’ CAG ACG TGT GCT CTT CCG ATC 3’) under the following conditions: 98 °C for 30 s, 15-17 cycles of 98 °C for 5 s, 65 °C for 10 s, with a final extension at 72 °C for 5 min. After the second PCR, the products were purified using the HP PCR product purification kit (Millipore Sigma), and eluted in 20 µL low TE (0.1X TE). For controls, the extracted plasmid DNAs were used as template: ~150 pg of the template was used for the first PCR using one of the HJ7210-7213 primers and one of the HJ7214-HJ7219 primers, both at 0.1 µM, with the similar conditions as for the cDNAs. Similarly, after column purification, 2 µL of the template from first PCR was used for the second PCR with similar conditions as for the cDNAs. All the products were analyzed by Agilent 2100 Bioanalyzer (Agilent Technologies, Santa Clara, CA) using high sensitivity DNA assay kit (Agilent Technologies) and the Qubit™ double stranded DNA Broad Range assay kit (Thermo Fisher Scientific) before pooling for library preparation.

| 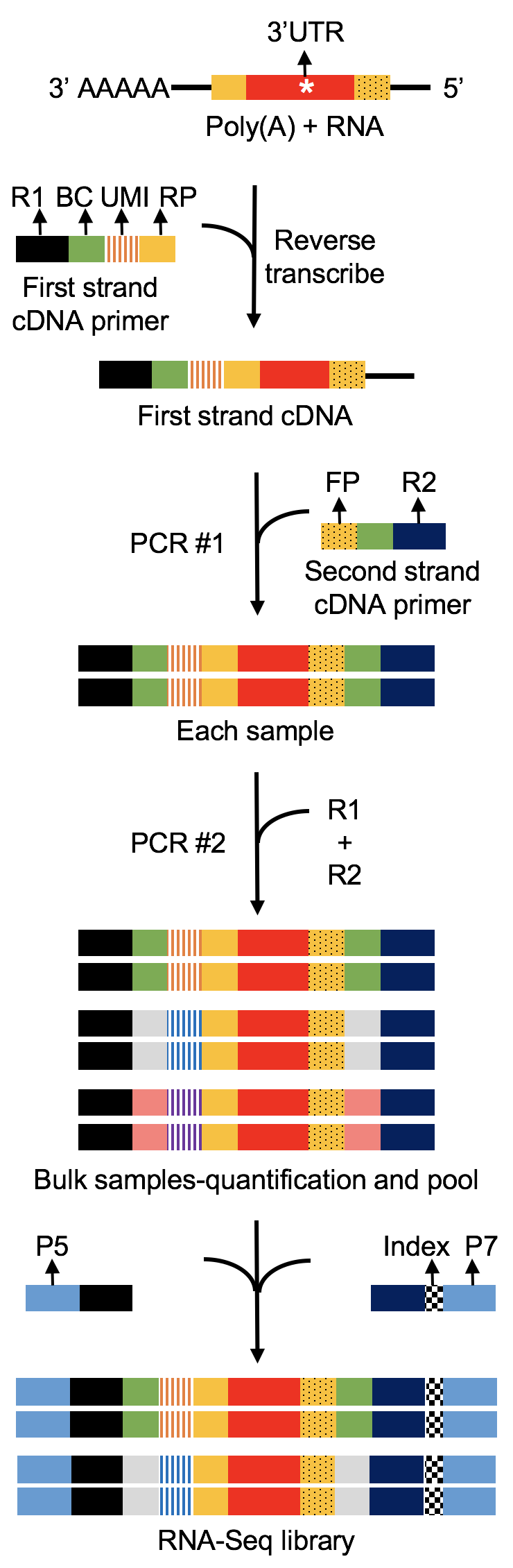 | **Figure S2. Modified library preparation for PASSPORT-seq.** First strand cDNA primers/ reverse transcription primers, HJ7210-7213 (R1=20 bp Read 1(partial Illumina adapter sequence), BC=9-12 bp sample barcode with staggered Ns, UMI=10 bp unique molecular indices, RP=22 bp universal reverse primer). Second strand cDNA primers, HJ7214-7219 (FP=22 bp universal forward primer, R2=21 bp Read 2 (partial Illumina adapter sequence). PCR amplification primers, HJ7220= 20 bp Read 1 (R1) and HJ7221= 21 bp Read 2 (R2). Library primers, Illumina i5 and i7 indexing primers were used to generate sequencing libraries. |
| --- | --- |

*Library preparation and sequencing*

The PCR fragments consisting of uniquely barcoded cDNAs and plasmid DNAs for each biological replicate were pooled together for each cell line (SH-SY5Y and SK-N-BE(2)). Next-generation sequencing libraries were prepared with 50 ng of PCR fragments in each reaction using Illumina TruSeq Nano DNA Low Throughput Library Prep Kit (FC-121-4001, Illumina, Inc., San Diego, CA). Because the PCR fragments already had adaptor sequences, DNA fragmentation, end-repair, dA-tailing, and adaptor ligation were omitted from the normal protocol. The PCR fragments were directly amplified using Illumina index adaptors (i5 and i7 indexing primers), using PCR conditions of 72 °C for 3 min, 98 °C for 30 s, followed by 5 cycles of 98 °C for 20 s, 60 °C for 15 s and 72 °C for 30 s, ending with 72 °C for 5 min. Each resulting library was quantified and the quality was accessed by Qubit™ double stranded DNA Broad Range assay kit (Thermo Fisher Scientific) and Agilent 2100 Bioanalyzer (Agilent Technologies), and both libraries were pooled in equal molarity. Five microliters of 3 nM pooled libraries, including 10% PhiX (Illumina), were then denatured, neutralized and applied to the cBot for flow cell deposition and cluster amplification before loading on to a lane of HiSeq 4000 (Illumina, Inc) for 75 nt paired-end sequencing. More than 360 million paired-end reads were generated, 96% of which reached Q30 (Phred quality score, 99.9% base call accuracy).

*Sequencing data analysis*

FASTQ files for DNA sequences were demultiplexed based upon the DNA barcodes identifying DNA source and replicate number using “cutadapt”, and the barcode and adapter sequences were trimmed from each read. The UMI was trimmed and stored in the read name using “umi_tools”. Using “bwa mem”, reads were mapped to chromosomes defined by the reference and alternative sequence for each SNP of interest. Finally, “umi_tools” was used to count the number of UMI-unique reads for each sequence. The overall workflow for data processing and analysis is depicted in Figure S3.

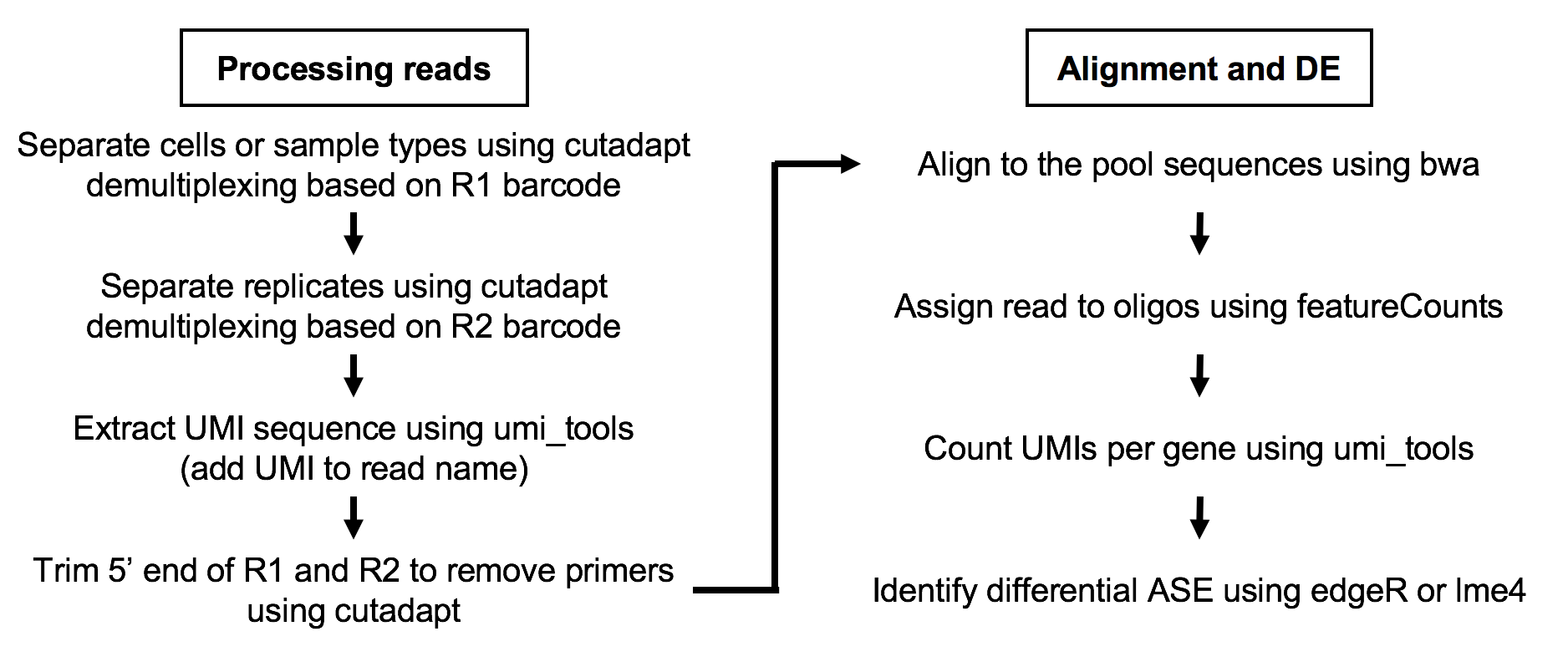

**Figure S3. Workflow for data processing and analysis pipeline for PASSPORT-seq.** Left panel, processing the reads, R1=Read1, R2=Read2. Right panel, sequencing processing and statistical model for identifying functional variants.

*Effect of ethanol treatment on SK-N-BE(2) cells*

To evaluate the potential effect of alcohol on the function of the 3’UTR SNPs, SK-N-BE(2) cells were grown to confluence, then treated with ethanol (10 mM and 20 mM). After 24 h and 42 h, cells were harvested for RNA isolation.

To count the reference and alternative alleles in ethanol-treatment RNA sequencing data, reads for each SNP of interest were extracted from the bam files using “bedtools intersect”. Then the reference and alternative alleles were counted using the “pysam” module of Python. A generalized linear mixed effect model was used to quantify the interaction between the alternative allele frequency of a SNP and ethanol treatment:

$$log(\mu)=\beta_{0}+\beta_{1}X_{1}+\beta_{2}X_{2}+\beta_{12}X_{1}X_{2}+bX_{S}+\varepsilon$$

where $\mu$ is the expected number of sequencing reads for one allele (reference or alternative), X_1_ is the allele type (0: reference allele, and 1: alternative allele), X_2_ is the subject study group (0: control group, 1: 10mM ethanol treatment group, and 2: 20mM ethanol treatment group), and X_S_ is the sample ID. In this model, $\beta_{0}$, $\beta_{1}$, $\beta_{2}$, and $\beta_{12}$ are fixed effect parameters, while b is a random effect that models the differences between study groups. In this model, the null hypothesis (H_0_) is: $\beta_{12}$= 0. Rejecting the null hypothesis indicates that the allelic imbalance differs between control and ethanol treatment group in a dose-response manner.
