## Supplemental Figures for "Allele-Specific Expression and High-Throughput Reporter Assay Reveal Functional Variants in Human Brains with Alcohol Use Disorders"

### Slide 1
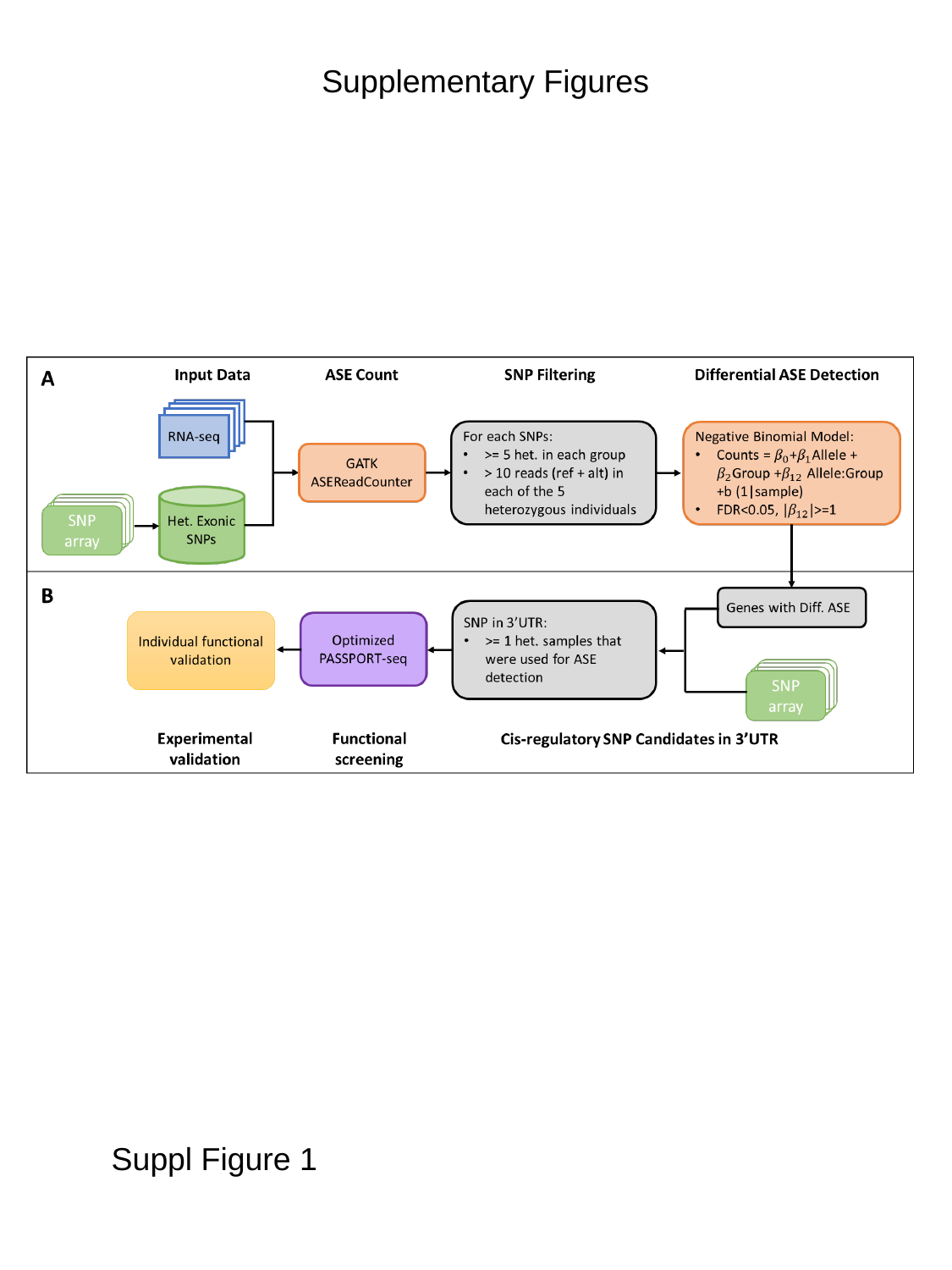

Supplementary Figures
Suppl Figure 1

### Slide 2
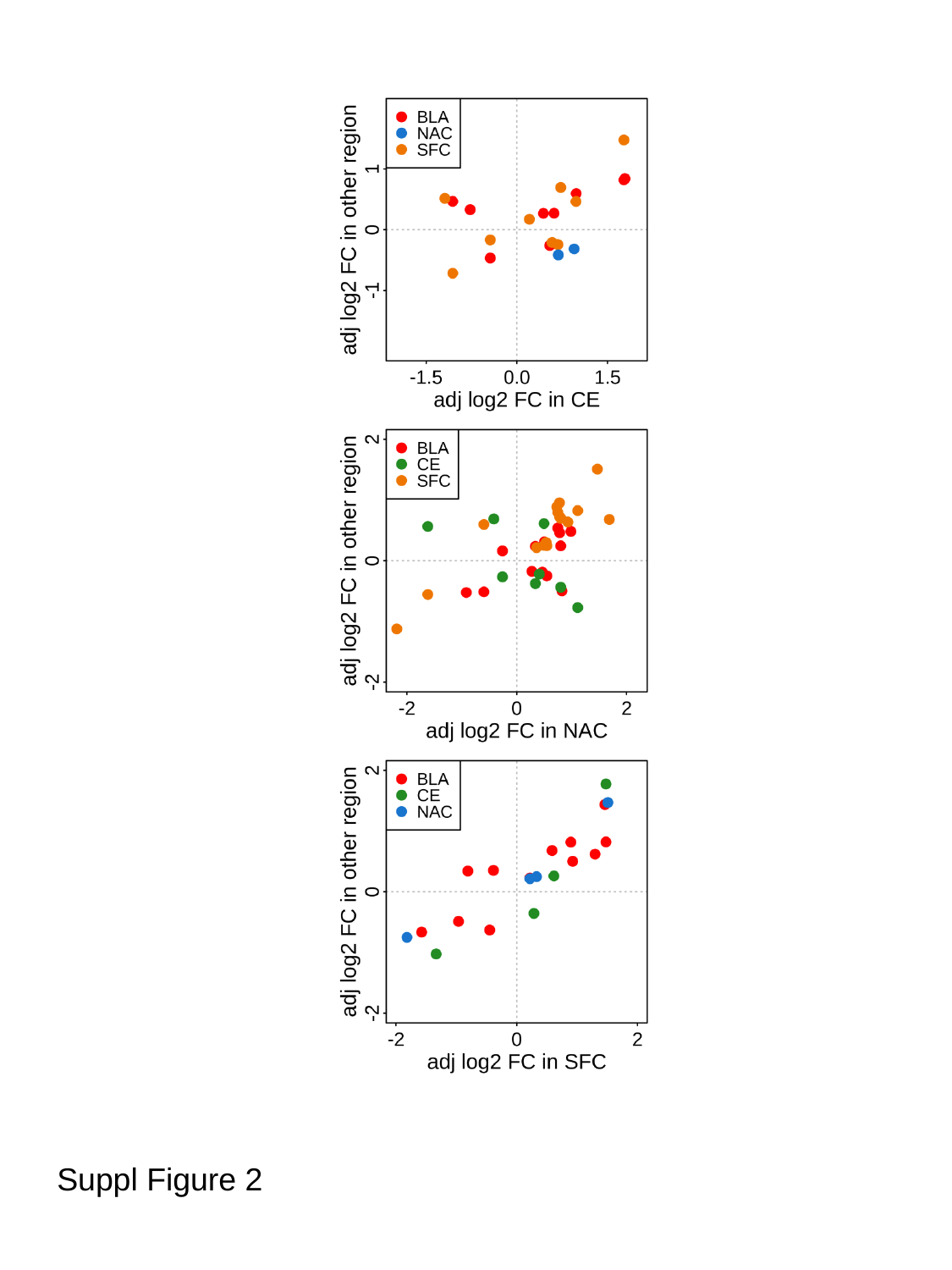

Suppl Figure 2

### Slide 3
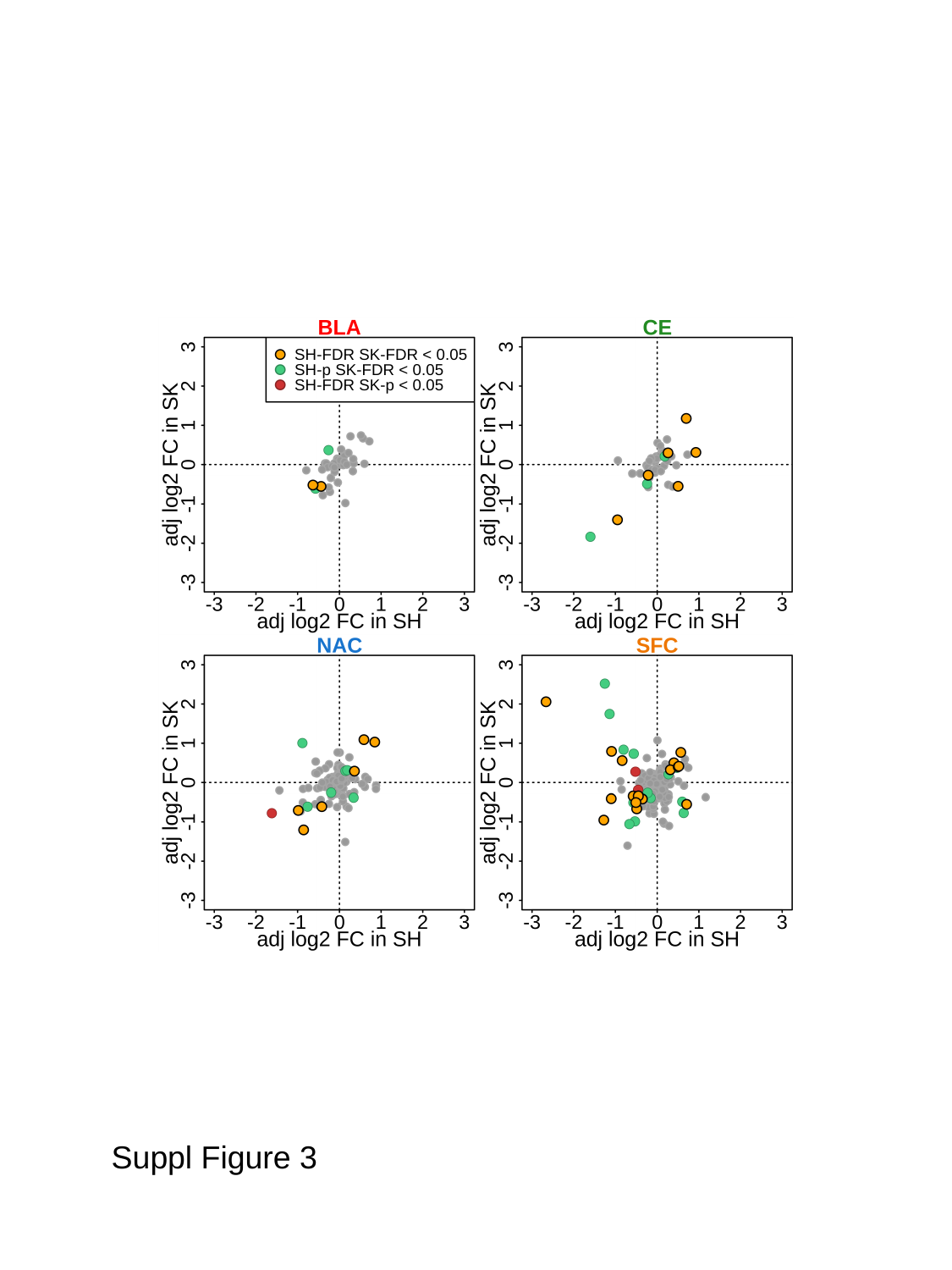

Suppl Figure 3

### Slide 4
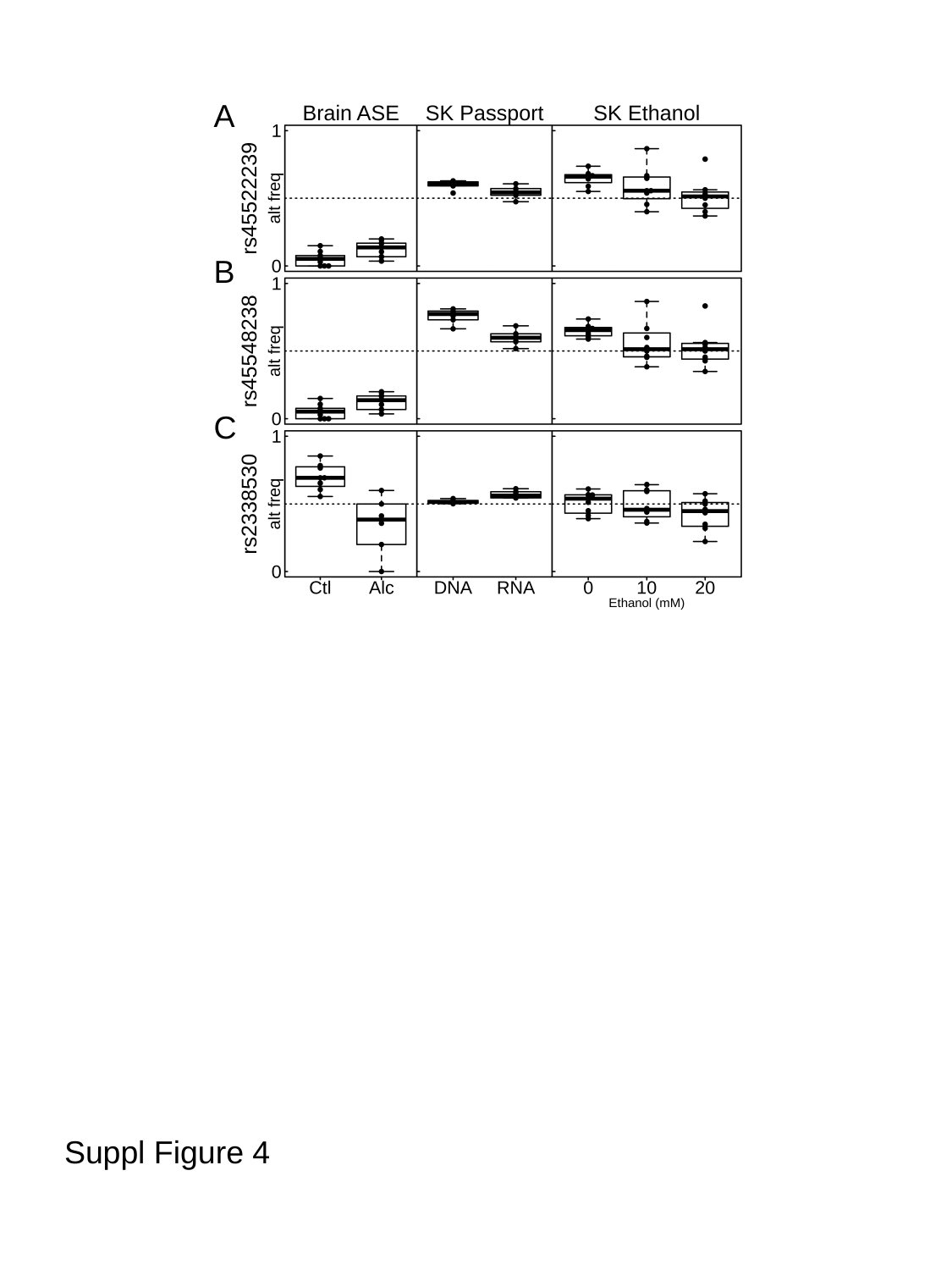

A
B
C
Suppl Figure 4
